# Behavioural diversity reveals distinct regimes of multisensory integration

**DOI:** 10.1101/2025.10.29.685323

**Authors:** Keon S. Allen, Douglas A. Ruff, Marlene R. Cohen

**Author notes:** Contributing authors.

## Abstract

Effective decision-making requires integrating multiple information sources, weighted by their reliability and context. While classic studies show near-optimal cue combination in extensively controlled laboratory settings and lengthy inperson experiments, everyday choices often occur in less controlled environments. We examined cue combination under these conditions using an online perceptual estimation task in large and diverse participant cohorts. Participants combined cues, including visual motion direction, spatial visual information, and sound. We investigated cue combination with and without cue conflict. Performance varied with age and self-reported ADHD or Autism. Visual cues were combined more optimally than audio-visual combinations. We observed qualitative similarities in an analagous task with non-human primates. We used electrical microstimulation in non-human primates, targeting unimodal or cross-modal association areas. Stimulation of visual cortex was integrated with sensory motion cues, while stimulation of prefrontal cortex promoted winner-take-all choices. These findings suggest distinctions between within- and across-modality integration, with deviations potentially informative of age or neurodiversity.

---

To navigate a complex and noisy world, humans must combine disparate sources of information. We dynamically integrate cues based on their reliability and contextual relevance. In perceptual decision-making, this process is referred to as cue combination, which is the integration of multiple sensory or feature-based signals into a single perceptual estimate. Cue combination has been a central focus of research on sensory integration and probabilistic inference [1, 2, 3].

Classic studies, conducted with lengthy in-person experiments in rigorously controlled laboratory environments, show that when cues are combined under ideal conditions, typically involving highly familiar signals, extensive training, or frequent feedback, observers often weight them in proportion to their reliability. This strategy produces judgments that approach statistically optimal estimates [4, 5, 2, 6].

Most natural decisions, however, occur in less controlled, more chaotic environments than laboratory settings. Here, we investigated how diverse participants performed in a brief, online version of a cue combination experiment with digitized visual and auditory cues. Further, we combined the large-scale behavioural testing in diverse human cohorts with causal manipulations in non-human primates to explore connections and generate hypotheses about how combination strategies emerge across modalities and individuals. Human participants performed an online perceptual estimation task involving combinations of visual and auditory cues. This design allowed us to assess integration in a way that approximates naturalistic decision-making. We compared performance across groups defined by age and self-reported neurodevelopmental diagnoses (attention deficit hyperactivity disorder, or ADHD, and Autism), revealing consistent differences in cue-weighting strategies [7, 8, 9].

For non-human primates, we paired visual motion stimuli with an artificial cue (electrical stimulation in either visual or prefrontal cortex) in non-human primates [10, 11, 12, 13, 14, 15, 16, 17, 18]. While stimulation of visual cortex was integrated with motion cues in a graded fashion, stimulation of prefrontal cortex resulted in winner-take-all (WTA) behaviour.

Our findings reveal circuit-level mechanisms underlying differences in cue combination within and across modalities and across groups, suggesting that the structure and function of cortical circuits involved in cue integration play a key role in shaping both behavioural strategies and individual differences in perceptual inference. More broadly, this work illustrates the value of combining large-scale human behavioural data with targeted neural interventions in animal models.

## Results

### A continuous localization task to estimate multisensory perception in a diverse set of participants

We begin by characterizing behavior in a continuous localization task designed to quantify both single-cue sensitivity and cue combination within the same individuals. This approach allows us to interpret multi-cue performance in relation to each participant’s own performance estimating each cue alone. By examining how performance changes when cues are combined within and across modalities, we establish a behavioral framework for testing alternative models of cue integration and for interpreting inter-individual variability.

We recruited a diverse set of human participants via an online platform (Prolific; Fig. 1A). Our participants consisted of 167 English-speaking participants (77 female, 80 male, 7 non-binary/gender queer, 3 transgender women) from around the world. Participants self-reported membership in one of four groups: 43 neurotypical younger adults (age range 18–35; mean age = 25), 43 neurotypical older adults (age range = 60–100; mean age = 64), 44 younger adults with ADHD (age range = 18–30; mean age = 25), and 37 younger adults with ASD (age range = 18–30; mean age = 26). All participants reported normal vision and hearing, and we verified headphone use during the task.

**Fig. 1.**
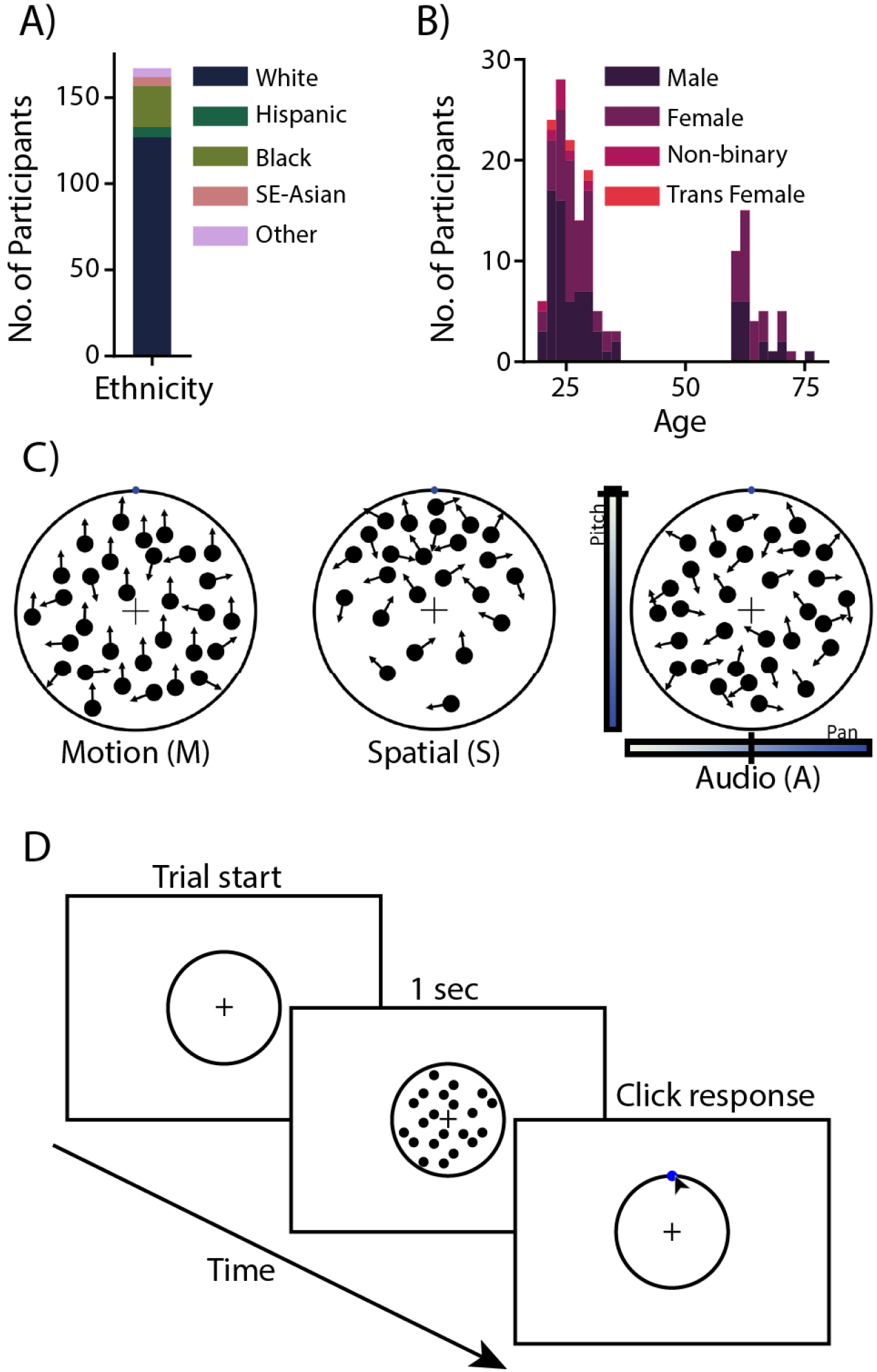
Human participants and task. A) Self-reported ethnicity of all 167 human participants. B) Self-reported gender and age of participants. Younger adults (restricted to ages 18-35) self-identified as having been diagnosed with ADHD, ASD, or neither. Older adults (restricted to ages 60+) self reported as not having been diagnosed with either ADHD or ASD. C) Experimental conditions. Each trial included one or two directional cues: the direction of motion of random dots (M), the spatial density of those dots (S), or an auditory cue whose pitch and pan indicated a direction (A). D) Schematic of an example trial. The trial began when a participant clicked on the central cross. Then the stimulus was displayed for 1 second, and the participant indicated a directional judgment by clicking the appropriate point on a target ring (blue point in this example was not shown).

We designed a continuous estimation task to measure participants’ perceptual and cue combination abilities. Participants reported, with a mouse click on the rim of a circle, the angle indicated by one or more visual or auditory cues (Fig. 1B). The visual cues were a random dot motion cue (M), in which a proportion of dots in the circle move coherently in one direction [19, 20] and a spatial cue (S), in which the dot locations were picked from a two-dimensional Gaussian whose center was at one angle around the circle. The auditory cue (A) was a simple pure tone whose pan and pitch indicated the target location. The pan of the sound was meant to evoke the sensation of interaural intensity differences, such that a tone which sounded louder in the left (right) ear indicated the target location was closer to the left (right) on the response circle. Similarly, as a proxy for sound elevation detection (which varies from person to person due to pinna shape), a tone that was higher (lower) in pitch indicated a location that was higher (lower) on the response circle. This two-dimensonal auditory cue allowed a simple sound to relate to any angle around the response circle. The audio cue was more intuitive than expected and participants were generally equally precise when using the audio cue as they were with the motion cue. Each block of trials involved one or more cues, the order of the blocks was randomized across participants, and the difficulty of the trials was identical for all participants. Participants received no feedback during the experiment except for a short tutorial, consisting of 8 practice trials at cardinal directions, preceding the single-cue conditions. The stimuli could indicate any angle around the circle, and when more than one cue was present, all cues indicated the same angle.

Participants typically clicked near the correct location, with modest error and little bias (Fig. 2A, B). Performance was generally better on multi-cue than on single-cue trials, as reflected in narrower response distributions (compare the width of the histograms in Fig. 2A for an example group). We quantified performance in each condition as the standard deviation of responses aligned to the correct location (e.g. standard deviation of the distributions in Fig. 2A), denoted *σ*.

**Fig. 2.**
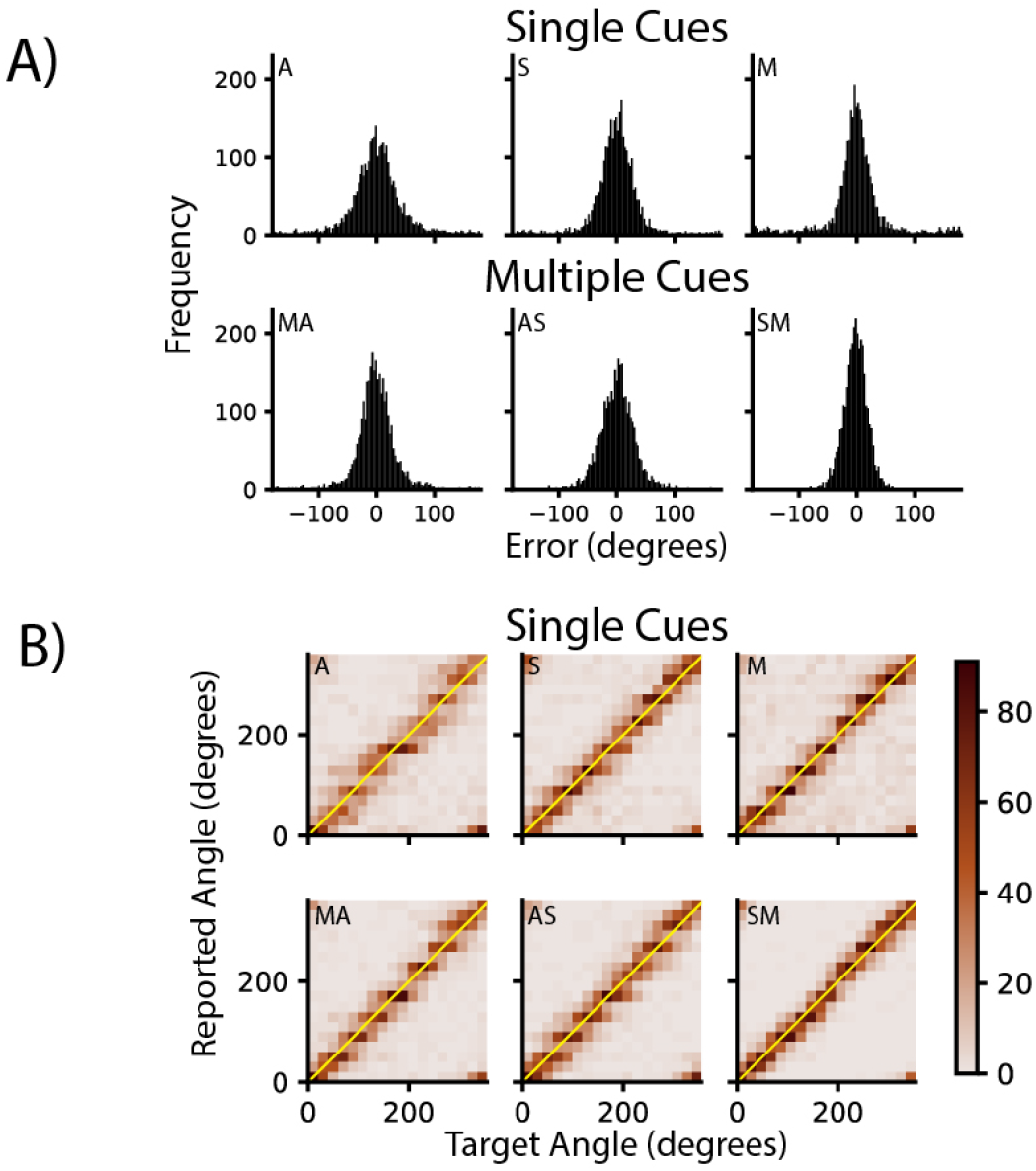
Choice patterns demonstrate that this example group of participants understood the cue combination and continuous nature of this task. A) Histograms, from the younger adult, non-ADHD, non-ASD group, of chosen direction relative to the actual direction for single-cue conditions (top row; M=motion, A=auditory; S=spatial) and trials containing two cues (bottom row; MA=motion-auditory, AS=auditory-spatial, and SM=spatial-motion). In general, multi-cue trials yield tighter distributions of choices, which follows the expectation that cue combination improves accuracy. B) Heatmaps of chosen versus actual direction for the same group and stimulus conditions as (A). Most choices are near the diagonal of these plots, indicating the lack of a prominent bias. The choices are distributed along the full range of choice directions, suggesting that the participant understood the continuous nature of the task. These results are typical of our data set.

We considered two alternative hypotheses for models to describe participant cue combination: the Bayesian optimal (OPT) model and the sub-optimal winner-take-all (WTA) model. If participants used a winner-take-all (WTA) strategy, selecting the cue they used most precisely in single-cue trials, their standard deviation in multi-cue trials would be:

*WTA*:

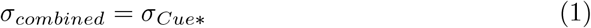

Alternatively, if participants combined cues according to a Bayesian optimal (OPT) strategy, weighting each cue by its reliability, the standard deviation in multi-cue trials would be:

*OPT*:

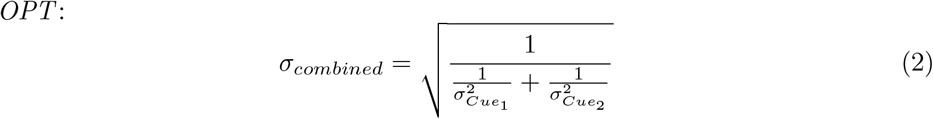

### Performance on single-cue trials varied by cue and group

Participants’ ability to use individual cues differed both across cues and participant groups (Fig. 3A). Although there was substantial variability across individuals, consistent patterns emerged when comparing visual motion, spatial, and auditory cues among younger and older neurotypical adults and younger adults with self-reported ADHD or ASD. There was substantial interaction between cue and group (two-way mixed ANOVA; F(15, 815) = 3.22, p<0.001), indicating that each group showed distinct sensitivities to different cues. In contrast, there was no overall effect of group [F(3, 163) = 1.62, p = 0.18], suggesting that these differences reflected selective strengths rather than general performance deficits.

**Fig. 3.**
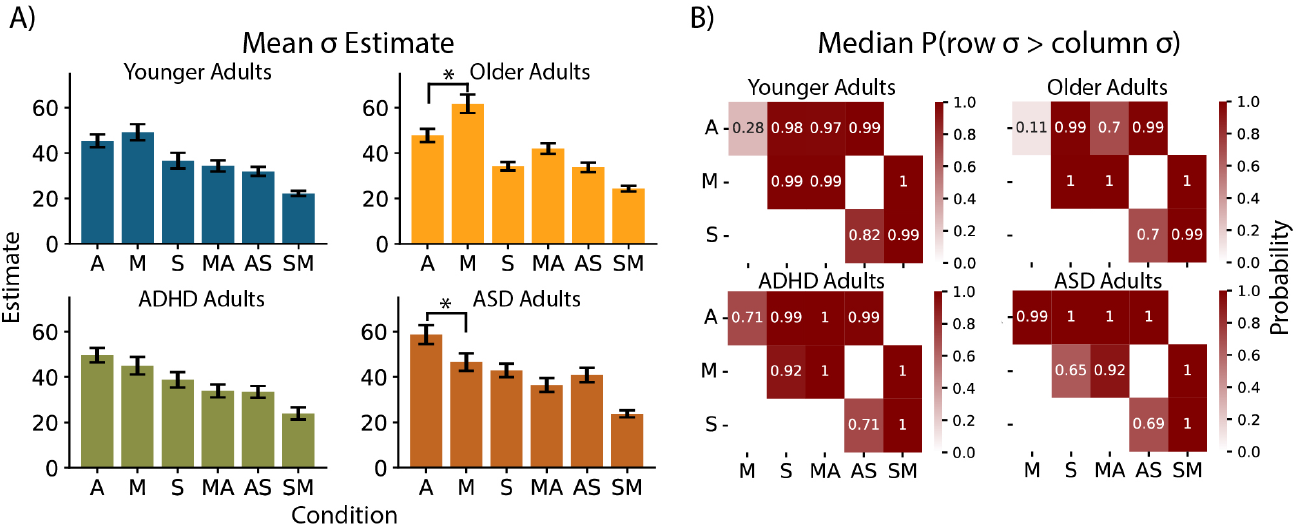
Participants performed better when they were able to combine information from multiple cues. A) Standard deviation of estimates for each condition (x-axes; A auditory, M motion, S spatial, and combinations of letters indicate multi-cue conditions) and for each group of participants (blue, neurotypical young adults; yellow, neurotypical older adults; brown, young adults with ASD; green, young adults with ADHD). Error bars represent standard error of the mean (SEM). B) Statistical comparisons of performance across each pair of conditions. Dark colors indicate that performance was better for the condition on the azimuth. (Numbers indicate the probability that the condition on the azimuth had better performance.)

Across all participants, performance differed significantly by cue (F(5, 815)= 67.10, p<0.001), but the specific pattern of strengths varied by group. Older adults performed more accurately on auditory than on motion trials [(t(42) = 2.99, p<0.01)], while younger adults with ASD performed better on motion than auditory trials [t(36) = 2.21, (p=0.03)]). There were no significant differences between participants with ADHD and age-matched neurotypical adults (Fig. 3A).

Together, these results show that sensitivity to single cues depends on both the cue and participant group. Importantly, these baseline differences underscore the need to interpret cue combination behaviour relative to each individual’s performance on single-cue trials. This consideration guided our subsequent analyses.

### Clear optimal integration for within-modality cues, but not for cross-modality cues

Participants combined information from multiple cues more effectively than from any single cue, as reflected by smaller response variability on multi-cue trials 3A, B), consistent with previous reports [4, 5, 10, 21, 22].

When combining two visual cues (motion and spatial), participants’ performance closely matched the optimal prediction. Their variability was significantly lower than predicted by the winner-take-all model, but not significantly different from the prediction of the optimal model. Fig. 4A, right). In contrast, when combining visual and auditory cues, performance was suboptimal, closer to the WTA prediction (Fig. 4A, left and middle).

**Fig. 4.**
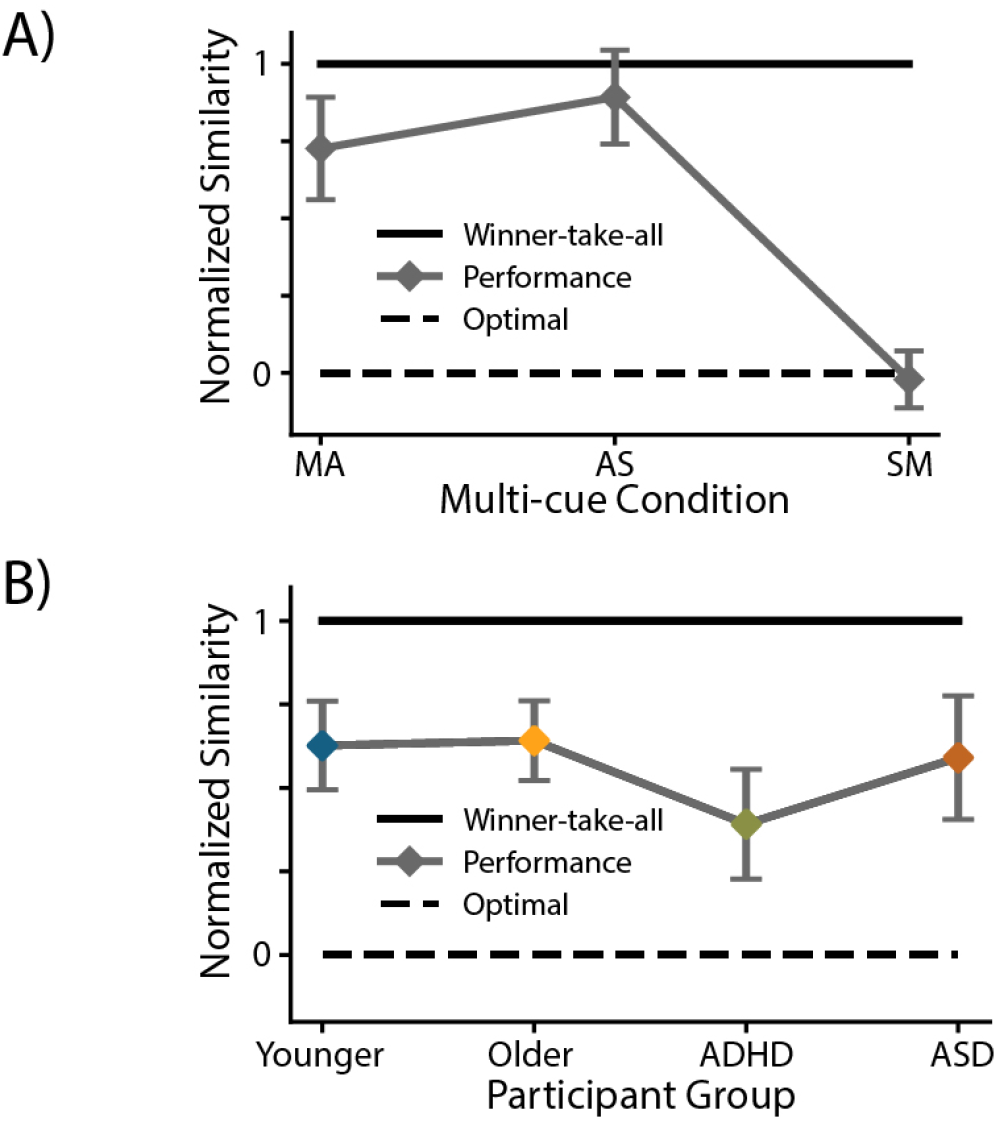
Cue combination was near optimal for two visual cues but suboptimal for visual-auditory combinations. A) Across all participants, split by condition, we quantified the similarity between performance on each multi-cue condition and the prediction of the winner-take all (solid line) and Bayesian optimal (dashed line) strategies as the normalized difference between optimal cue combination performance and observed performance in the better of the two single cues and the relevant two cue condition. Error bars represent SEM, and data are averaged across all participants in all groups. Performance on the cross-modal conditions (MA and AS) supports a winner-take-all model: the difference between the winner-take-all prediction and combined condition performance was not significantly different for the MA [(t(166) = 0.82, p>0.05)] and AS [t(166) = 1.48, p>0.05)] cross-modal conditions. In contrast, performance on those conditions was significantly different from the prediction of the optimal model for the MA [(t(166) = 6.61, p<0.001)] and AS [t(166) = 10.86, p<0.001)] conditions. Performance on the within modality condition supports a Bayesian optimal model: performance on SM trials was significantly different from the prediction of the winner-take-all model [t(166) = 1.48, p<0.001)] but was not significantly different from the prediction of the Bayesian optimal model [t(166) = 0.26, p=0.80)]. (SM) supports Bayesian optimal performance. B) The normalized similarity between the performance of each group of participants and the predictions of the two models, averaged over conditions, was not significantly different across groups (p>0.05).

Some evidence for a distinction between within and across modality cue combination exists within neurophysiological and psychophysical observations. Visual cues are often combined within the visual system itself [23], and this combination occurs even in anesthetized animals [24]. Multisensory cue combination, however, likely involves computations in association areas that may require additional processing or effort [25, 26]. These studies show that within-modality cue combination is less demanding in multi-cue learning than multisensory cues, perhaps suggesting differences in how effectively or effortfully those cues are combined.

Across participant groups, cue combination performance were broadly similar. Averaged across all multi-cue conditions, all four groups showed performance intermediate between the WTA and optimal predictions (Fig. 4B).

### Cue conflict experiments reveal group differences

The cue combination experiment generated clear predictions about how participants in different groups would weight information from different sensory cues. We tested these predictions in a cue conflict experiment in which the directions indicated by the two cues on multi-cue trials were offset by 15° in opposite directions from the target angle (Fig.5).

**Fig. 5.**
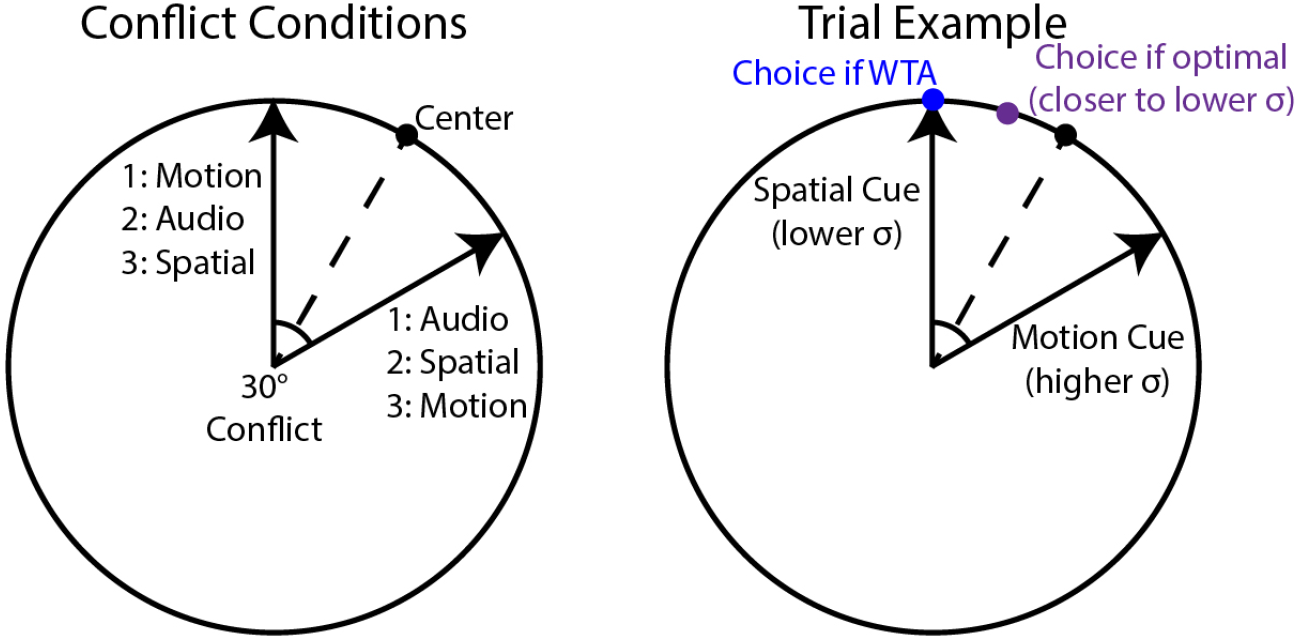
Conditions in the cue conflict experiment. Each multi-cue condition included an offset of 30° between the two cues (e.g. in the motion-auditory condition, the motion direction was always 30° counter-clockwise of the direction indicated by the auditory cue). On an example spatial-motion trial (right), the winner-take all model predicts that participants should select the direction of the single cue on which the participant performed best (e.g. spatial in this example; blue dot), while the optimal model predicts an intermediate choice biased toward the direction of the better single cue (purple dot).

Although there was considerable individual variability, participants’ PSEs were generally shifted toward the location of the cue on which they performed better on single-cue trials (Fig. 6). We quantified this by comparing the actual PSE (grey triangles in Fig. 6; colored triangles show group means) with the prediction of the Bayesian optimal model (black markers in Fig. 6).

**Fig. 6.**
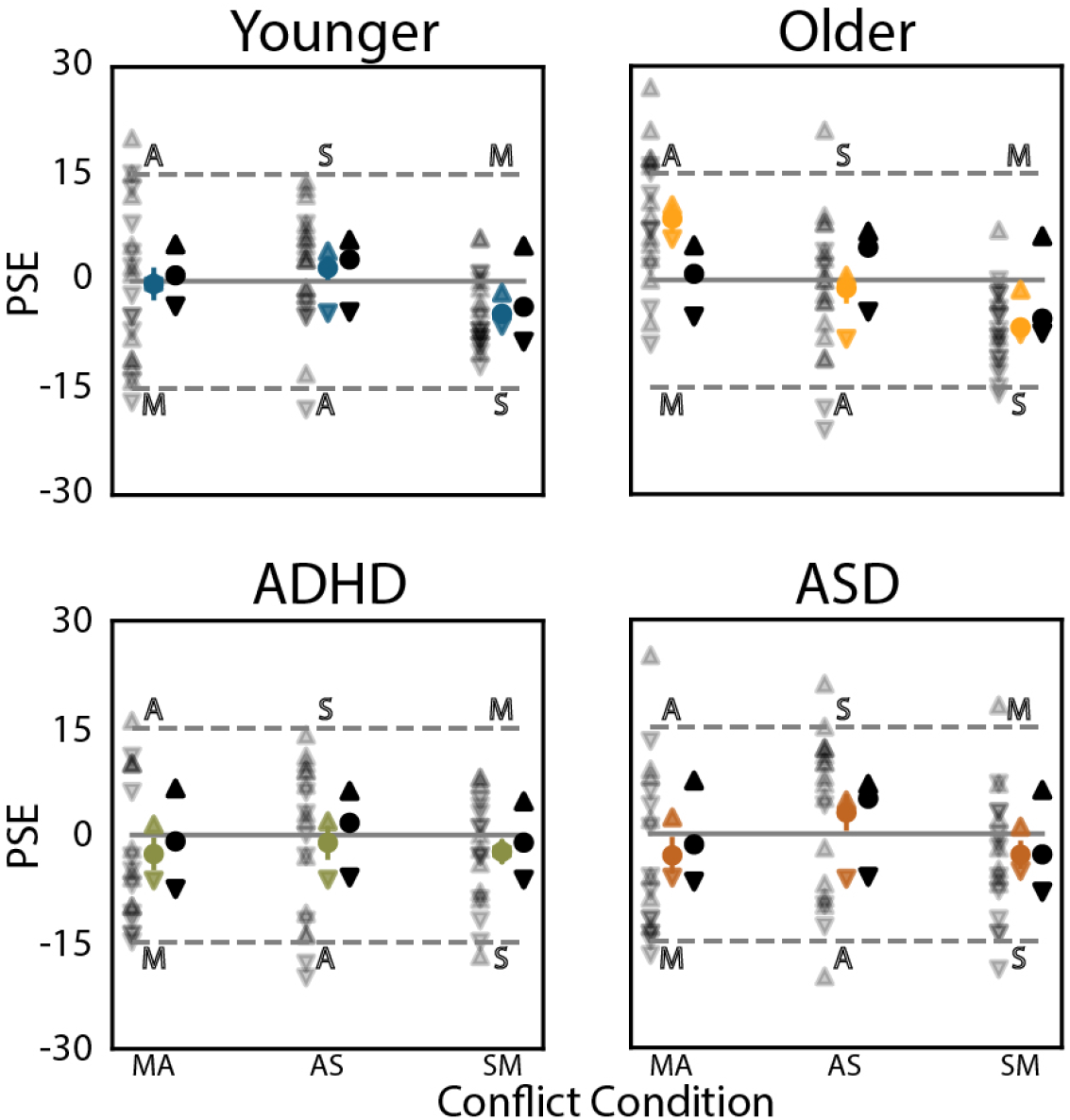
Points of subjective equality (PSEs) for each group of participants, for each conflict condition. The scattered gray triangles show data for each participant. The up-pointing (down-pointing) triangles represent participants for whom, of the pair of cues involved in the conflict, the cue on the upper (lower) dashed line was better. The solid, colored markers show the means for participants. The solid black markers show the mean predictions according to the optimal model.

One consistent exception was observed in older adults, who relied more heavily on the auditory cue than on either visual cue (compare the yellow circle to the black marker in (Fig. 6)), even when it was suboptimal to do so. This pattern persisted even among older adults who performed better with motion or spatial cues on single-cue trials (compare the triangles to the black circle in Fig. 6)).

### Distinct integration strategies for signals arising in sensory and association cortex

The behavioral findings of sub-optimality during cross-modal cue combination motivated complementary experiments in non-human primates designed to probe underlying mechanisms. We trained two rhesus monkeys (Macaca mulatta, both male, 8 and 10 kg) to perform an analogous eye movement version of the motion estimation task performed by our human participants (Fig. 7A; different aspects of these data have been published previously [27]. While fixating a central point, the monkeys viewed a random dot motion stimulus that could move in any direction and at any coherence level. They reported their perceived motion direction by making an eye movement to a corresponding location on a circular target ring. The amount of fruit juice reward on each trial depended on the accuracy of their estimate (see Methods). For reasons unrelated to the current study, some choices were associated with higher rewards, which produced a choice bias [27]. The monkeys’ choices closely tracked the actual motion direction, biased by expected reward (most points near or slightly below the diagonal in Fig. 7B).

**Fig. 7.**
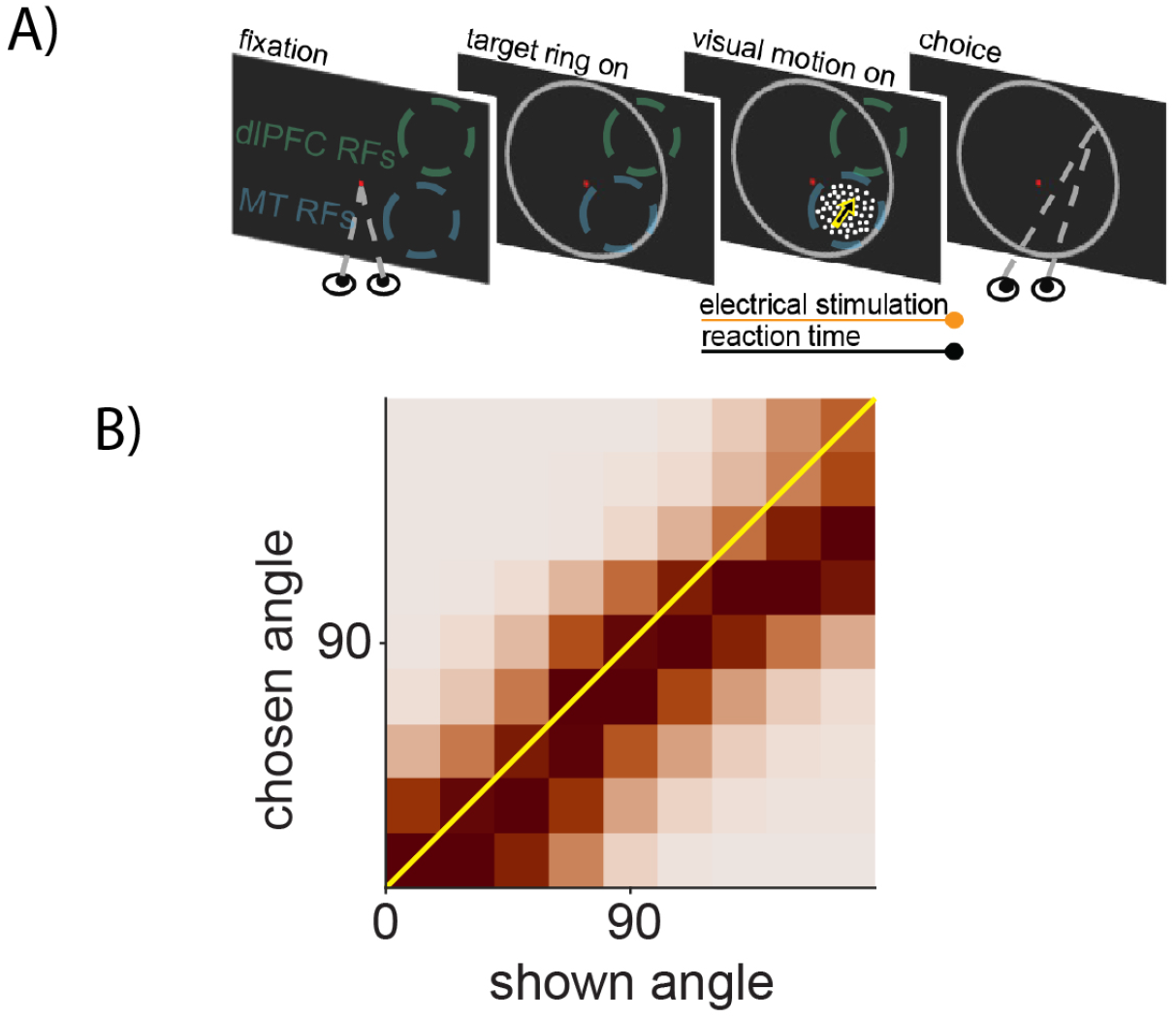
Causal experiment to compare cue combination within the visual system and in association areas. A) Continuous estimation task for our monkey microstimulation experiment. Monkeys fixated a central spot and viewed a random dot motion stimulus that, like in our human experiments, could move in any direction. Whenever they were ready, the monkeys indicated their choice by looking at a point on the target ring that corresponded to a direction judgment. The dots disappeared at the moment the monkeys initiated the eye movement. The monkeys were rewarded according to the accuracy of their motion direction estimate, which was sometimes scaled according to a rule that is irrelevant for the current study (see Methods). During the experiments, we recorded populations of neurons in MT and dlPFC using multielectrode probes. We placed the dots within the joint spatial receptive fields of the recorded MT neurons and positioned the target ring so that a portion of it overlapped the response fields of the recorded dlPFC neurons. On a randomly selected subset of trials, we electrically microstimulated one electrode at a time in MT or dlPFC during the period that the dots were displayed. B) Composite of behavioural data across 29 sessions from 2 monkeys during non-microstimulation trials. In general, the monkeys achieved high accuracy and their choices varied smoothly with the motion direction.

While the monkeys performed this task, we recorded from groups of neurons in the middle temporal area (MT), a region critical for motion perception [28, 29, 20], and the dorsolateral prefrontal cortex (dlPFC), an association area involved in multisensory integration, decision-making, reward processing, and eye movement planning [30, 31, 32]. As in previous work, [17, 20], we then perturbed activity in these areas using low-current electrical microstimulation (20–40 *mu*A) applied to a single electrode during stimulus presentation on randomly interleaved trials. This approach allowed us to test how information introduced at different cortical levels was combined with visual motion signals during perceptual decisions.

Although the monkey and human experiments share a common computational goal—estimating motion direction under uncertainty, they differ in important respects, including task structure, training history, and the nature of the signals being combined. Accordingly, these experiments were not designed to establish cross-species equivalence in cue combination behavior. In particular, electrical microstimulation is not intended to mimic a natural sensory cue, and differences between microstimulation and auditory or visual signals are unavoidable. Instead, the purpose of the primate experiments is to leverage a key advantage unavailable in human studies: the ability to control the cortical source of a decision-related signal. By introducing an artificial signal at different stages of the cortical hierarchy, we can directly test how information originating in sensory versus association areas is integrated with visual evidence during perceptual decisions. This causal manipulation allows us to move beyond behavioral dissociations and ask how the site of signal generation shapes the strategy by which information is combined.

Consistent with the prediction from human psychophysics, visual motion and electrical stimulation of MT were combined in a near-optimal way. As in previous studies [20], microstimulation in MT biased choices toward the preferred direction of the neurons near the stimulated site, producing intermediate estimates between the visual and electrical cues. To test whether this reflected Bayesian optimal weighting, we varied the coherence of the visual motion stimulus, thereby changing its reliability. The Bayesian optimal model predicts that stimulation should have a stronger influence on choices when the visual cue is less reliable. Our MT microstimulation data matched this prediction: the bias toward the stimulated direction was greater when motion coherence was low than when it was high (compare rows in Fig. 8).

**Fig. 8.**
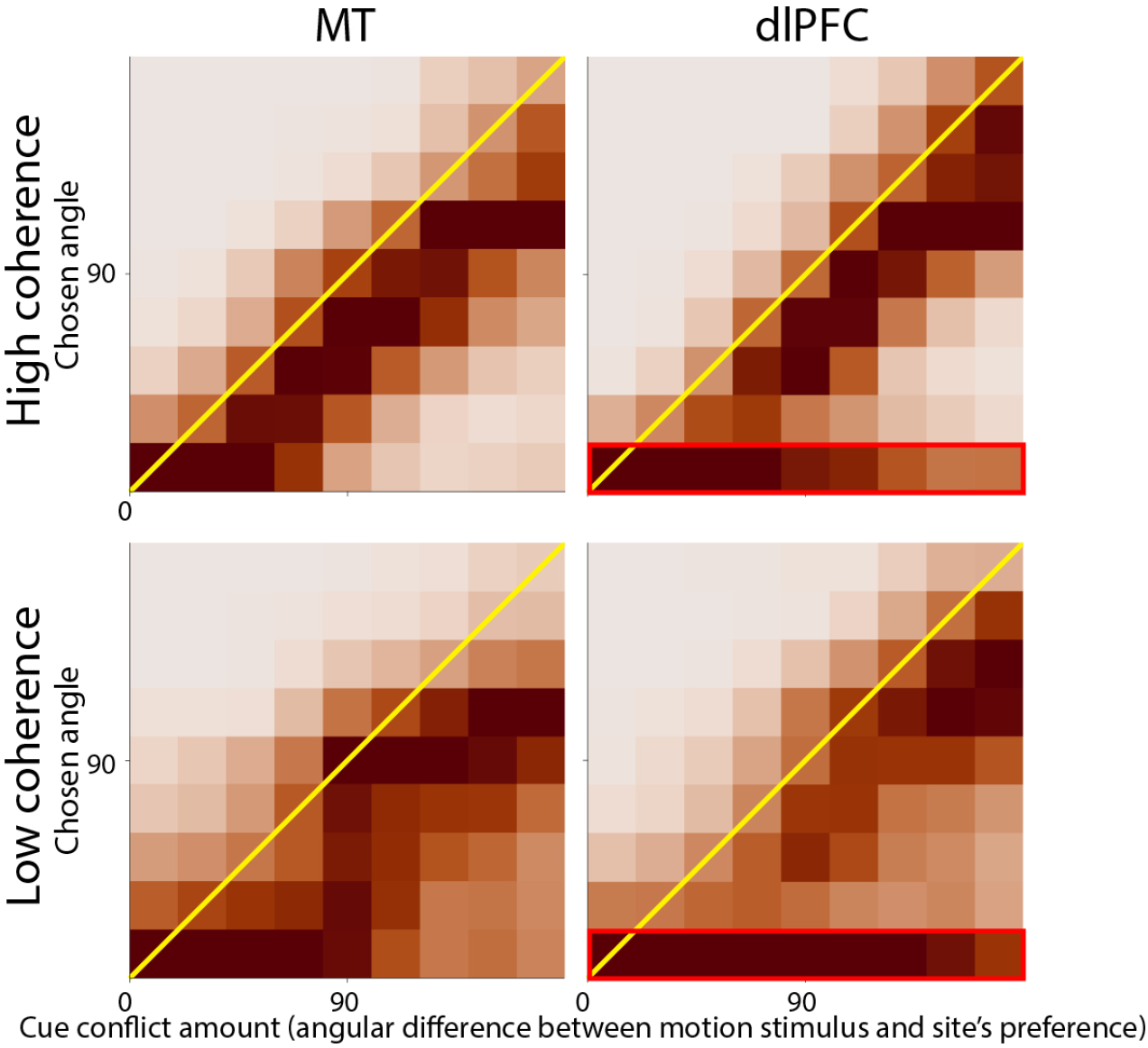
Causal evidence that cue combination within the visual system (e.g. area MT) matches predictions of optimality while being suboptimal in association areas (e.g. dlPFC). The heat maps depict choices from trials where we stimulated area MT(left) or dlPFC(right), when the visual stimulus had high (top) or low (bottom) coherence. MT microstimulation biased choices toward the preferred direction of the stimulated neurons while dlPFC stimulation produced bimodal choices that were directed either to the preference of the stimulated neurons or to the mean choice during non-stimulation trials (horizontal bands, highlighted with red border). Hartigan’s Dip tests reveal both sets of dlPFC choices are significantly bimodal (p<.001) and the MT ones are not (p>.05). Decreasing the coherence increased the bias on MT microstimulation trials and increased the proportion of choices toward the preference of the stimulated neurons, but did not change the unimodal or bimodal nature of the distributions.

In contrast, stimulation of dlPFC produced a sub-optimal pattern similar to the cross-modal cue combination observed in humans on our online task. The model predicts that stimulation of dlPFC should produce bimodal choice distributions, with one mode aligned with the visual motion direction and the other aligned with the preferred direction of the stimulated neurons. Moreover, the proportion of trials in each mode should depend on the relative reliability of each cue. Consistent with this prediction, dlPFC microstimulation produced bimodal choice distributions centered on the visual and electrical cues, and the proportion of stimulation-driven choices increased when the visual stimulus was less reliable (low coherence) compared with when it was more reliable (high coherence; Fig. 8).

## Discussion

We combined behavioural, physiological, and causal approaches across species, sensory modalities, ages, and neurodiversities to investigate the neuronal computations that underlie decisions based on multiple sources of information. The human and non-human primate results, taken together, suggest circuit level hypotheses for the observed variability in within- and across-modality cue combination. Mechanisms supporting cue combination can depend on whether information is integrated within a single sensory system or across modalities, and may vary across individuals, particularly with age and neurodiversity.

### Complementary benefits of laboratory neurophysiological studies and online psychophysics experiments

Our approach integrated neurophysiological recordings and perturbations in monkeys with online psychophysical testing in humans, two methodologies that offer complementary strengths. Neurophysiological experiments provide exceptional control over behaviour and the ability to record and manipulate neuronal activity. These experiments have revealed that single neurons can combine multiple sources of information in a way that reflects the reliability of each cue, suggesting a plausible substrate for cue combination [10]. However, these studies are necessarily limited in the number of subjects and therefore cannot easily address questions about age, diagnosis, or individual variability.

Online psychophysics provides the opposite advantage. By recruiting large, diverse cohorts of human participants, it is possible to examine variability across individuals and groups at an unprecedented scale. Although online testing has less experimental control than laboratory studies—for example, participants self-report demographic information and eye position cannot be tracked—the behaviour we observed was lawful and internally consistent.

In our study, the human results generated a neural hypothesis that we were able to test directly in non-human primates. The convergence between these two approaches illustrates how controlled animal neurophysiology and large-scale human psychophysics can be used together to identify general computational principles of perception.

### Age-related differences in multisensory cue combination

We found consistent differences between how older and younger adults used and combined cues to guide decisions. Unlike younger adults, older adults performed better on auditory than motion trials (Fig. 3A) and tended to over-rely on auditory cues during combined motion–auditory trials. Even older adults who performed better on motion than auditory cues alone gave disproportionate weight to auditory information in the cue conflict experiment (Fig. 6), a pattern not observed in younger adults.

Several age-related changes in perception and cognition may contribute to this suboptimal cue combination. Older adults often experience declines in the reliability of sensory inputs, such as reduced visual acuity and auditory sensitivity, leading to noisier perceptual representations [33]. Both younger and older adults can benefit from multisensory integration in simple detection or visuo-haptic tasks [34, 35, 36], although in some contexts, older adults show diminished multisensory benefits ([37, 38]. Audiovisual integration has also been observed to be altered in older adults, compared to younger adults. [39] Increased susceptibility to multisensory illusions, such as the McGurk effect, also suggests that older adults may have greater difficulty estimating the reliability of conflicting cues [40].

Age-related declines in attentional control may also affect cue weighting by making it more difficult to flexibly allocate attention across modalities[33]. These results highlight that changes in multisensory integration across the lifespan may reflect both sensory degradation and alterations in cognitive control processes.

### Implications for neurodiversity in multisensory integration

Individuals with ASD performed well overall but showed selective difficulty with auditory cues. Individuals with ASD may be less susceptible to multisensory illusions [41, 9, 42], potentially reflecting reduced multisensory integration, especially of conflicting information [43]. Other studies, however, have found intact integration for visual–vestibular cues [44] or for visual navigation tasks, albeit with greater variability [45]. Our results align with this latter view, showing that multisensory integration in ASD may be preserved but cue-specific.

Participants with ADHD performed at least as well as the other groups, including neurotypical younger adults. Future studies could investigate the role of certain features of ADHD, such as heightened responsiveness to salient or novel information, on integration. Studying situations in which individuals with ASD or ADHD perform as well as, or better than, neurotypical participants can identify mechanisms of resilience and inform new approaches to intervention or accommodation.

## Methods

### Cue combination and conflict tasks for human participants

We built our online human psychophysics experiments using Gorilla experiment building tools [46] and customized scripts. We recruited adult human participants through Prolific (prolific.co), who performed our cue combination task (Fig. 1). All procedures involving human participants were approved by the Institutional Review Board of the University of Chicago (and granted a Category 4 exemption).

167 English-speaking participants (77 female, 80 male, 7 non-binary/gender queer, 3 transgender women) from around the world participated in one of two experiments. We did not observe substantially different results in male and female participants. Participants self-reported as belonging to one of the following groups: 43 younger adults (mean age: 25), 43 older adults (mean age: 64), 44 ADHD adults (mean age: 25), 37 ASD adults (mean age: 26). All participants reported normal vision and hearing and completed the task wearing headphones.

Before beginning the experiment, participants had to succeed a headphone test. During a given trial in the test, white noise would be played in multiple intervals. On one interval, a quiet tone would be played simultaneously. Participants would report which interval contained the masked sound. Sufficient performance on this test would be difficult without the use of headphones rather than sound speakers.

On a each trial, clicked a mouse on the rim of a circle to report the direction indicated by one or more visual or auditory cues. The visual cues were a motion direction cue, in which a proportion of dots within the circle move coherently in one direction, and a spatial cue, in which the dot density was higher in one part of the circle. The audio cue was a simple tone where the pitch and pan of the sound indicated a location on the circle.

The auditory and visual stimuli were generated using Python 3 and relevant packages (numpy, matplotlib, pydub, ffmpeg, etc.). Each stimulus was presented as a short one-second video clip at 60 frames per second and a resolution of 800 by 800 pixels. On a given trial, a circle appeared on screen, the inside of which was populated with smaller, moving dots. On the first frame of the video, each dot was generated with a random ‘age’ up to the maximum ‘life’ of 30 frames. Upon reaching the maximum lifespan, a dot would be removed and replotted in a new location according to the parameters of the condition. In the Motion (M) condition, the dots were uniformly distributed within the circle. On a given frame, each dot would move one step in the direction of either the target angle or a uniformly random angle around the circle. The proportion of dots moving at the target angle on any given frame is the coherence, which was set to a value of 0.4. Thus, in order to perform the task, participants could only rely on motion information. In the Spatial (S) condition, the coherence of the dots was set to 0 and on any given frame, each dot would move in a random direction. Instead, dots within the circle were plotted according to a two-dimensional gaussian centered on the target angle on the rim of the circle. As such, the dots subtly clustered near the target angle and to perform the task, participants would need to make use of the spatial organization cue. In the Audio (A) condition, a given auditory stimulus was a simple tone (sine wave) played for 1 second. The tones mapped onto locations around the circle with the horizontal component of the angle mapping onto auditory pan ranging from -1 (left only) to 1 (right only) and the vertical component mapping onto frequencies ranging from 400Hz to 800Hz, well within the hearing range for younger and older adults. This was a general, low-tech mimicry of auditory location direction that avoided considerations such as head and pinna shape while leveraging existing expectations such that it was both intuitive and informative. On screen, the dots within the circle had a coherence of 0 and were uniformly distributed within the circle. Thus, in order to perform the task, participants had to rely on the audio cue alone.

Combined conditions were pairwise combinations of the three individual cues. The spatial and motion (SM) combined condition was parameterized the same way as in the motion only condition except, rather than dots plotted according to a uniform distribution, they were plotted according to the spatial organization of the spatial condition. The audio and spatial (AS) combined condition and the audio and motion (AM) combined condition were the same as the respective spatial-alone and motion-alone condition with the appropriate audio playing alongside the video.

These were uploaded as videos to the Gorilla.sc experiment platform to build the experiment. Single-cue conditions and multi-cue conditions were counterbalanced separately, with single-cue conditions preceding multi-cue conditions in the experiment. This was necessary to minimize task confusion for participants. Preceding each single-cue condition were tutorial videos and 8 practice trials to ensure participants understood the task. Tutorial videos were created in Adobe After Effects, animating a mouse over the stimuli to demonstrate how the participant should give their response on a given trial. The 8 practice trials that followed occurred at cardinal directions around the circle with corrective feedback.

In the perceptual estimation experiment, the presented cues were always consistent with one another. That is, the audio, motion, and spatial cues all indicated the same location on the rim of the circle.

Each experimental condition utilized one or more of the motion (M), spatial (S), or audio (A) cues.

The cue conflict task was the same as above, except the multi cue conditions were symmetrically offset in opposite directions by 15° each from the target location.

### Behavioural task for rhesus monkeys

The subjects for the monkey experiments were two adult rhesus monkeys, S and O (Macaca mulatta, both male, 8 and 10 kg, respectively). Different analyses of a subset of these data have been published previously [27]. Before behavioural training, the monkeys were implanted with a custom titanium head holding device. All surgical and behavioural procedures conformed to the guidelines established by the National Institutes of Health and were approved by the Institutional Animal Care and Use Committees of the University of Pittsburgh and Carnegie Mellon.

During behavioural training and experiments, the monkeys sat in a primate chair that provided head restraint. We presented visual stimuli on a high-quality LCD monitor (VPixx Technologies Inc., QC Canada; 120Hz refresh rate) positioned 57cm from the monkey’s eyes. We monitored eye movements and eye position using an infrared optical eye tracker (Eyelink 1000, SR Research, ON, Canada). Behavioural control and stimulus presentation were managed by computers using a Unix based operating system (running MATLAB, The MathWorks, Inc., Natick, MA, and PsychToolbox [47]).

The monkeys were trained to perform a variant on the continuous direction estimation task performed by our human participants, but there was only a motion cue. A trial began when the monkey fixated a central spot on a black background (inside an invisible 1° diameter window). Then a circular target ring was displayed and remained visible for the remainder of the trial. After a randomly selected delay (range 200-400 ms), a dynamic random dot kinematogram was displayed in a stationary circular aperture inside the target ring. During these multi-coherence experiments, trials using two motion coherences were randomly interleaved (low coherence range: 15% - 20%; high coherence range: 35% - 50%).

Monkeys were trained to make a saccadic eye movement to a location on the target ring to signify their choice. The monkeys could initiate the eye movement between 150 and 1500 ms after visual stimulus onset. The reward was based on the accuracy of the motion estimate. For reasons unrelated to the current study and described in [27], the reward associated with some choices was scaled by an amount indicated by the location of pink- or purple-colored portions of the target ring. The data presented in Fig. 8 are only from the randomly interleaved half of trials in which high reward expectation was associated with the preferred direction of the stimulated neurons.

### Electrophysiological methods

Before electrophysiological recordings and microstimulation experiments, two recording cylinders (Crist instruments Co., Inc., Hagerstown, MD) were placed over the principal sulcus and visual cortex to provide access to dlPFC and MT, respectively. We recorded from groups of neurons in MT and/or dlPFC using 24 or 32 channel linear probes (V- and S-probes; Plexon Inc, Dallas, TX) positioned using grids (Crist Instruments Company Inc., Hagerstown, MD) and advanced using a hydraulic microdrive (Kopf instruments, Tujunga, CA). Spiking activity, local field potentials, eye position, and task events were recorded at 30,000 samples/second (using Trellis software and Ripple recording hardware; Ripple, Salt Lake City, UT). All data were analyzed using custom scripts (MATLAB; The MathWorks, Inc., Natick, MA or Python).

Details of our recording methods have been reported previously [27]. Briefly, MT units were identified based on stereotactic coordinates, gray-white matter transitions, and functional properties. In most experiments, the receptive fields of the recorded MT units were less than 12 degrees eccentric so we could place the motion stimulus within their receptive fields without overlapping the target ring. We measured the spatial receptive fields and motion direction tuning to verify electrode positioning, guide motion stimulus positioning, and identify the tuning preferences of sites recorded on the contacts we used for electrical stimulation.

We recorded dlPFC units along the banks and gyri of the principal sulcus (determined by stereotactic location and visual inspection during surgery). We included recordings from locations in which we observed spatial selectivity during any period of a delayed saccade task. We typically recorded units whose response fields were more than 10 degrees eccentric so we could position a portion of the target ring within them and avoid overlap with the visual motion stimulus.

For electrical microstimulation, we selected two contacts on a probe to be stimulated on random subsets of trials (at most one contact was stimulated on each trial, and we also interleaved no stimulation trials). Microstimulation experiments were only ever performed in one brain area per session. The MT units on candidate stimulation contacts were required to have a suitable visual receptive field location and strong direction tuning. The dlPFC units on candidate stimulation contacts were required to have suitably positioned visual or premotor response fields.

Electrical microstimulation began with the onset of the visual motion stimulus and remained on until the monkey moved its eyes out of the fixation window to initiate the eye movement to indicate its motion direction judgment. Microstimulation was biphasic and delivered at 200 Hz with an amplitude that ranged across sessions between 20 and 40 mA (8-10).

## Data Analysis

The location of each click (humans) or eye movement (monkeys) to indicate a decision was stored in x, y coordinates. Subsequent analyses are based on the angle between the clicked location and the target (correct) location.

### Bayesian Parameter Estimation

Many analyses are based on estimates of the mean (*µ*) and standard deviation (*σ*) of choices. We estimate these from discrete data by calculating the likelihood of each possible *µ, σ* pairing using Equation 3 where *θ*_*i*_ is the calculated angle for a given trial.

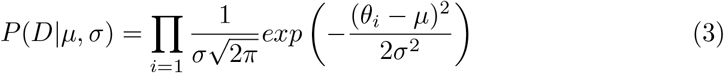

Using the likelihoods with a uniform prior, we calculated joint posterior distributions over *µ, σ* using Bayes’ formula (Equation 4) with the prior term having been canceled out. Marginalizing over *µ*, we get a probability density over *σ*. The mode of this PDF is our *σ* estimate.

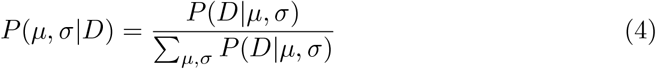

Joint posterior probabilities are calculated the same way for the conflict experiment. The *σ* estimates were calculated by marginalizing the joint posterior over *µ* and it was marginalized over *σ* to get the the *µ* estimates which is also the point of subjective equality (PSE).

### Bayesian Comparison of *σ* Distributions

We use the *σ* probability density functions to compare the single and multi-cue conditions to one another, calculating the probability that one PDF is greater than another. These results were summarized in Figure 3B. We let *ρ*_*row*_ stand for the *σ* PDF pertaining to a given condition and *ρ*_*col*_ stand for the *σ* PDF of another condition. Equation 5, where *x*_*i*_(*x*_*j*_) refers to the *i*th(*j*th) bin of the PDF, describes the calculation.

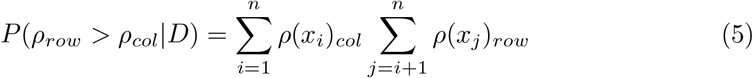

### Cue Combination Predictions

For a given participant, we took 10,000 samples from their single-cue *σ* PDFs. For each combined condition, the relevant single-cue samples were combined according to the different model equations (1, 2). This resulted in PDFs over *σ* for each model. The modes of these PDFs were taken as the predictions of each model. These data were summarized in Figure 4 where median performance was plotted normalized between the predictions of the winner-take-all and optimal models.

### Frequentist Statistics

The two-way mixed ANOVA and subsequent post-hoc tests were performed using the most likely *σ* values (maximum a posteriori estimates) obtained from the earlier parameter estimation. We used publicly available Python packages including *pandas, scipy, pingouin*, and *statsmodels*.

## Acknowledgements

We thank Karen McCracken for technical assistance with the animal experiments and Ramanujan Srinath, Lily Kramer, Sylvia Durian, Grace DiRisio, Cheng Xue, Kursti Ropp, and other members of the Cohen lab for helpful discussions and comments on an earlier version of the manuscript. This work was supported by the Simons Foundation (Simons Collaboration on the Global Brain award 542961SPI to M.R.C), and the National Institutes of Health (awards R01EY022930, R01EY034723, and RF1NS121913 to M.R.C).

## Declarations

The authors declare no competing financial interests. Data and code will be provided upon reasonable request.

## Notes

### Competing Interest Statement

The authors have declared no competing interest.

### Summary of Updates

Edits to text and figures for clarity.

